# Whole-brain dynamics and hormonal fluctuations across the menstrual cycle: The role of progesterone and age in healthy women

**DOI:** 10.1101/2023.07.23.550200

**Authors:** Daniela S. Avila-Varela, Esmeralda Hidalgo-Lopez, Paulina Clara Dagnino, Irene Acero-Pousa, Elvira del Agua, Gustavo Deco, Belinda Pletzer, Anira Escrichs

**Affiliations:** Computational Neuroscience Group, Center for Brain and Cognition, Department of Information and Communication Technologies, Universitat Pompeu Fabra, Barcelona, Catalonia, Spain; Linguistic Center (CLUL), School of Arts and Humanities, University of Lisbon, Lisbon, Portugal; Department of Psychology & Centre for Cognitive Neuroscience, Paris-Lodron-University Salzburg, Salzburg, Austria; Department of Psychology, University of Michigan, Ann Arbor, MI, USA; Chronic Pain and Fatigue Research Center, Department of Anesthesiology, University of Michigan, Ann Arbor, MI, USA; Institució Catalana de la Recerca i Estudis Avançats (ICREA), Barcelona, Catalonia, Spain

**Keywords:** menstrual cycle, whole-brain dynamics, progesterone, estradiol, resting-state fMRI, metastability, integration, dynamical complexity

## Abstract

Recent neuroimaging research suggests that female sex hormone fluctuations modulate brain activity. Nevertheless, how brain network dynamics change across the female menstrual cycle remains largely unknown. Here, we investigated the dynamical complexity underlying three menstrual cycle phases (i.e., early follicular, pre-ovulatory, and mid-luteal) in 60 healthy naturally-cycling women scanned using resting-state fMRI. Our results revealed that the preovulatory phase exhibited the highest variability over time (node-metastability) across the whole-brain functional network compared to the early follicular and mid-luteal phases, while the early follicular showed the lowest. Additionally, we found that large-scale resting-state networks reconfigure along the menstrual cycle phases. Finally, we used multilevel mixed-effects models to examine the impact of hormonal fluctuations and age on whole-brain and resting-state networks. We found significant age-related changes across the whole brain, control, and dorsolateral attention networks. Additionally, we observed progesterone-related changes, specifically within limbic and somatomotor networks. Overall, these findings evidence that both age and progesterone modulate brain network dynamics along the menstrual cycle.

## Introduction

Approximately 49.7% of the world’s female population are women of reproductive age (15 to 45 years old). Of these women, around 58% experience a natural menstrual cycle (free of hormonal contraceptives), characterized by physiological fluctuations of the ovarian hormones estradiol and progesterone (Dubol et al., 2021; Roos et al., 2015; United Nations, 2019a,b). A regular menstrual cycle ranges between 21 to 35 days with less than seven days of variability in cycle length (Fehring et al., 2006) and can be divided into three main phases according to the hormonal levels. The onset of menses initiates the follicular phase and marks the beginning of the menstrual cycle. The follicular phase is characterized by low progesterone levels and a gradual increase in estradiol concentrations until the pre-ovulatory phase, in which estradiol levels reach their peak. Following ovulation, the luteal phase commences and continues until the last day of the menstrual cycle. This phase is characterized by an increase in progesterone levels, which reach their highest peak in the middle of the phase (Bull et al., 2019; Reed and Carr, 2000). Converging and growing evidence shows that these cyclic hormonal fluctuations impact brain structure and function in multiple ways (Arélin et al., 2015; Barth et al., 2016; Dubol et al., 2021; Fitzgerald et al., 2020; Mueller et al., 2021; Pritschet et al., 2020, 2021; Rehbein et al., 2021; Taylor et al., 2020). Especially, large-scale network dynamics have proven responsive to endogenous hormonal fluctuations (Hidalgo-Lopez et al., 2021; Pritschet et al., 2020).

Resting-state fMRI enables exploring the brain’s intrinsic organization of large-scale distributed networks. Recent neuroimaging studies on the menstrual cycle have revealed cycle-related brain activity fluctuations in several resting-state networks, including the default mode network (DMN), salience, dorsal attention, and subcortical networks (Engman et al., 2018; Hidalgo-Lopez et al., 2020; Petersen et al., 2014; Pletzer et al., 2016; Pritschet et al., 2020; Weis et al., 2019). The DMN connectivity has been consistently reported to change across the cycle phases. For instance, studies have reported increased functional connectivity to the dorsolateral prefrontal cortex in late follicular (Weis et al., 2019) while decreased functional connectivity to bilateral angular gyri (Hidalgo-Lopez et al., 2020; Petersen et al., 2014) and increased connectivity to the cuneus during the luteal phase (Pletzer et al., 2016). Dynamic changes across the cycle phases have been shown to involve a reorganization of salience and executive control networks depending on the hormonal levels (Hidalgo-Lopez and Pletzer, 2021). During the luteal phase, higher functional connectivity of salience and limbic networks (Engman et al., 2018; Hidalgo-Lopez et al., 2020; Pletzer et al., 2016) has been reported, but also oscillatory activity and functional connectivity of subcortical areas (Hidalgo-Lopez et al., 2020; Pletzer et al., 2016).

To date, only a few menstrual cycle fMRI studies have used a whole-brain dynamic approach and showed that progesterone and estradiol impact brain dynamics and information processing across large-scale brain networks (De Filippi et al., 2021; Greenwell et al., 2023; Mueller et al., 2021; Pritschet et al., 2020). Such an approach can provide a deeper understanding of how hormonal fluctuations modulate the underlying brain dynamics. Nonetheless, the literature on the menstrual cycle and brain dynamics is limited and shows inconsistent findings, which have been attributed to variations in methodological approaches, the lack of a standard definition of cycle phases and small sample sizes (Engman et al., 2018; Hidalgo-Lopez et al., 2020; Schmalenberger et al., 2021). Addressing these limitations is critical for providing a more reliable understanding of how menstrual cycle-related changes impact cognition, emotion, and behavior, as well as for developing targeted interventions for menstrual cycle-related disorders.

The dynamic framework proposed by Deco and Kringelbach (2017); Deco et al. (2017b) provides a comprehensive view of the brain as a complex system that exhibits specific dynamical properties essential for effective information processing. This framework emphasizes the concept of intrinsic ignition, which refers to the ability of brain areas to ignite and sustain neural activity that can propagate across the whole-brain network by measuring metastability (Deco et al., 2017a). Metastability refers to the capacity of the brain to flexibly engage and disengage different brain areas over time, reflecting the variability of global synchronization. Higher metastability corresponds to more complex brain dynamics, while reduced metastability points to more stable dynamics. This framework has proven remarkably robust in capturing subtle differences in whole-brain dynamics across several brain states in health and disease, including deep sleep, meditation, aging, depression, and abnormal development, among others (Deco et al., 2017b; Escrichs et al., 2021, 2019; Mayneris-Perxachs et al., 2022; Padilla et al., 2020). Understanding the underlying whole-brain dynamics is crucial for examining the effects of factors that impact brain function, such as menstrual cycle phases and hormone fluctuations.

In this study, we aimed to investigate the dynamical complexity of the menstrual cycle phases by examining resting state activity at different levels of analysis, including global, network and local brain activity patterns in a sample of 60 healthy naturally-cycling women scanned using resting state fMRI. Specifically, we computed the intrinsic ignition framework (Deco and Kringelbach, 2017) across the whole-brain network and within eight well-known resting-state networks (control, DMN, dorsolateral attention, limbic, somatomotor, salience, subcortical, and visual) along three menstrual cycle phases (early follicular, pre-ovulatory, and mid-luteal). Furthermore, we employed multilevel modeling to examine the effects of hormone fluctuations (progesterone and estradiol) and age on brain network dynamics. We hypothesize that hormone fluctuations and age can substantially influence the metastability of whole-brain and resting-state network dynamics.

## Methods

### Participants

A total sample of 60 healthy young women was selected from a dataset previously described in Hidalgo-Lopez et al. (2021). The study was conducted on women between the ages of 18 and 35 who had regular menstrual cycles lasting 21 to 35 days with intercycle variability of fewer than 7 days. All participants met the following inclusion criteria: (1) not having used hormonal contraceptives in the 6 months before the study, (2) having no history of neurological, psychiatric, or endocrine disorders, and (3) not taking any medications. Three appointments were scheduled for each participant: during the early follicular phase (1-7 days after the onset of current menses), in the pre-ovulatory phase (2-3 days before the expected date of ovulation), and during the mid-luteal phase (3 days after ovulation to 3 days before the expected onset of next menses). The order of appointments was counterbalanced to minimize any potential bias. The pre-ovulatory sessions were confirmed by commercial urinary ovulation tests (Pregnafix ®). Participants were asked to confirm the onset of the following menses after the appointment. The cycle duration was calculated based on the participants’ self-reported onset dates of their last three periods. Participants gave informed written consent to participate in the study. The University of Salzburg’s ethics committee approved the study and conformed to the Code of Ethics by the World Medical Association (Declaration of Helsinki).

### Hormone analysis

Hormone levels were assessed using Salimetrics salivary Estradiol and Progesterone ELISAs. Saliva samples were collected from participants using the passive drool method, stored at -20° until analysis, and centrifuged twice at 3000 rpm for 15 and 10 min, respectively, to remove solid particles. All samples were tested in duplicates, and samples with more than 25% variation between duplicates were re-analyzed. To study the effects of the menstrual cycle on estradiol and progesterone levels, a linear mixed model was fitted using estradiol and progesterone as dependent variables, cycle phase as fixed effects, and subject number as a random effect. The analysis was performed in R 4.2.2 using the package lme4 (Bates et al., 2015). To calculate pairwise comparisons between cycle phases, we used the multcomp package (Hothorn et al., 2008); and *p*-values were adjusted for multiple comparisons with the Bonferroni method, where we divide the standard significance level by the number of tests performed (*i*.*e*., .05*/*3 = .017).

### MRI data acquisition

MRI images were acquired using a Siemens Magnetom TIM Trio 3T scanner. The high-resolution T1-weighted images were acquired with 160 sagittal slices (slice thickness=1 mm; TE=291 ms; TR=2300 ms; TI delay=900 ms; (FA) 9°; FOV 256X256 mm). Resting-state fMRI was performed using a T2* weighted gradient echo-planar (EPI) sequence with 36 transversal slices (TE=30 ms; TR=2250 ms; flip angle (FA) 70°; slice thickness=3.0 mm; matrix 192×192; FOV 192 mm; in-plane resolution 2.6×2.6 mm). During the resting state, participants were instructed to relax, close their eyes, and let their minds flow.

### Preprocessing

For each participant, the first six volumes were discarded, and the functional images were despiked using the 3d-despiking algorithm implemented in AFNI. Then, despiked images were pre-processed using standard procedures and templates in SPM12 (www.fil.ion.ucl.ac.uk/spm), including segmentation of the structural images using CAT12. The resulting images were then subjected to the ICA-AROMA algorithm implemented in FSL to remove artefactual components in a non-aggressive manner. Finally, BOLD time series were filtered within the narrowband (0.01–0.09 Hz) and extracted according to a resting state cortical atlas of 100 nodes and 7 networks (Schaefer et al., 2018), and a subcortical atlas of 16 nodes (here referred to as the subcortical network) (Tian et al., 2020).

### Intrinsic ignition framework

We computed the Intrinsic Ignition Framework (Deco and Kringelbach, 2017) to study the dynamical complexity across the whole brain functional network and within eight well-known resting-state networks across three menstrual cycle phases (early follicular, pre-ovulatory, and mid-luteal). This framework quantifies the degree of whole-brain integration from spontaneously occurring events over time. **Figure 1** schematizes the methodology for obtaining the intrinsic integration across brain areas. The algorithm captures driving events for each brain area, which are transformed into a binary signal using a threshold (Tagliazucchi et al., 2012). To represent events as a binary signal, time series are transformed into z-scores, denoted as *z*_*i*_(t), and a threshold value, *θ*, is applied. Specifically, an event is marked as 1 in the binary sequence *σ*(t) if *z*_*i*_(t) exceeds the threshold from below and marked as 0 otherwise. When a brain area triggers an event, neural activity is measured in all brain areas within a set time window of 4TR. A binary matrix is then constructed to represent the connectivity between brain areas exhibiting simultaneous activity. The measure of global integration (Deco et al., 2015) is applied to determine the broadness of communication across the network for each driving event (i.e., the largest subcomponent). This process is repeated for each spontaneous neural event to obtain the node-metastability (measured as the standard deviation of the integration over time) for each brain area across the network, where higher node-metastability corresponds to higher dynamical complexity.

**Figure 1:**
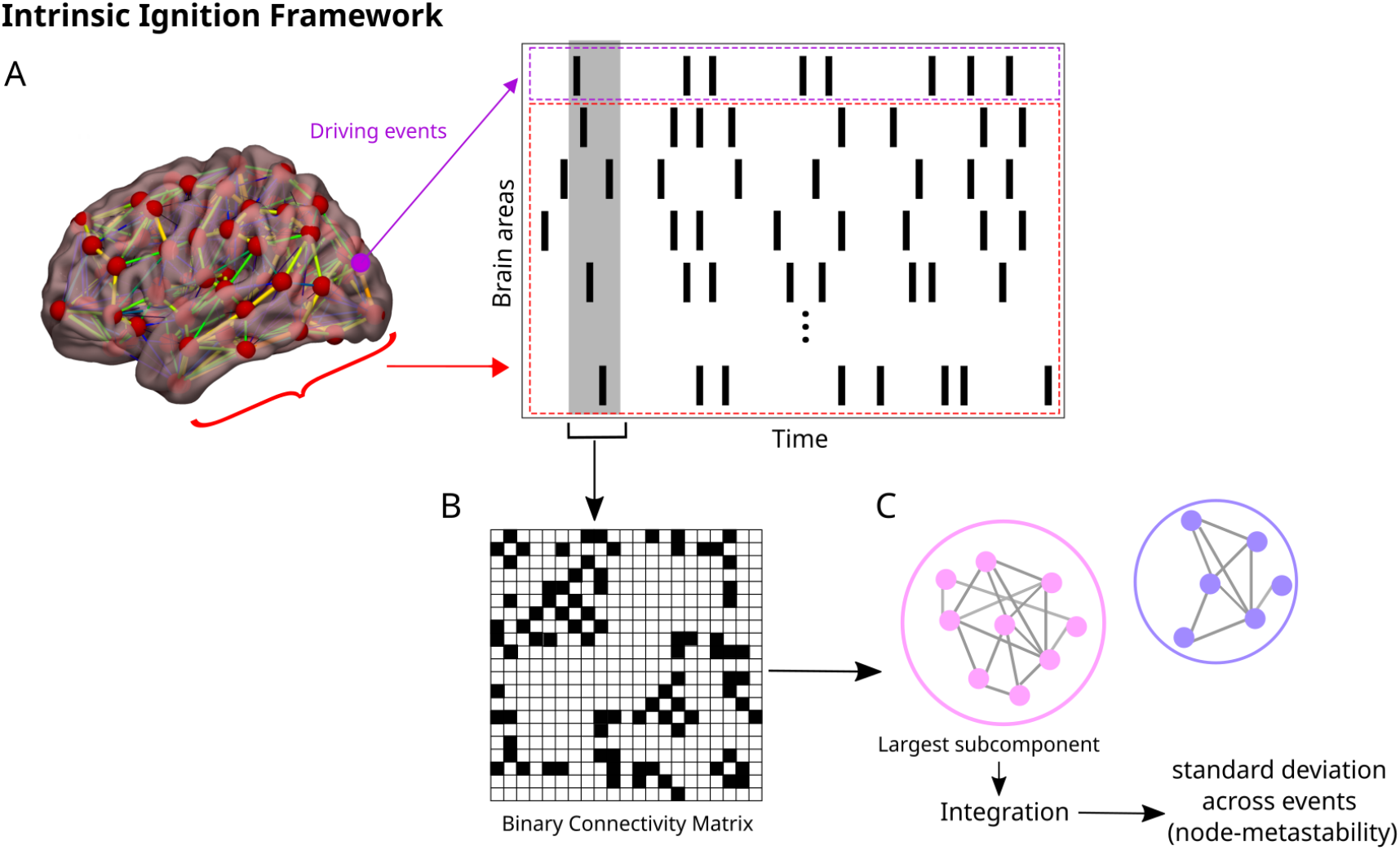
Measuring intrinsic ignition. **(A)** Events were captured applying a threshold method (Tagliazucchi et al., 2012) (see purple area). For each event elicited (gray area), the activity in the rest of the network was measured in the time window of 4TR (see red area). **(B)** A binarized matrix was obtained, representing the connectivity between brain areas where activity was simultaneous. **(C)** Applying the global integration measure (Deco et al., 2015), we obtained the largest subcomponent. Repeating the process for each driving event, we calculated the node-metastability computed as the standard deviation of the integration of each brain area over time.

### Statistical analyses

We applied the Monte Carlo permutation method (10,000 iterations) to evaluate the outcomes of the node-metastability between each pair of phases to be compared. Furthermore, we employed the False Discovery Rate (FDR) method (Hochberg and Benjamini, 1990) to account for multiple comparisons when analyzing the differences between phases across the whole brain network and within the eight resting-state networks. Moreover, we carried out mixed-effect multilevel models. In particular, we included the metastability values as the dependent variable (the whole brain network and for each resting-state network, respectively), fixed effects (age, progesterone, estradiol, and the interaction between progesterone and estradiol) and the random effects (subject, age, progesterone and estradiol).The resulting model syntax is *Metastability ∼* 1 + *age* + *estradiol* + *progesterone* + *estradiol ∗ progesterone* + (1|*estradiol*) + (1|*progesterone*) + (1|*age*) + (1|*sub*).

## Results

### Demographic data

This study analyzed a cohort of 60 young healthy women with a regular menstrual cycle as previously reported in (Hidalgo-Lopez et al., 2021). The participants had an average age of 25.4 years (range 18-35 years), and their average cycle length was 28 days (range 23-38). Estradiol levels were significantly higher during the pre-ovulatory than the early follicular (*Estimate* = 0.329, *SE* = 0.051, *p <* 0.001) or mid-luteal phase (*Estimate* = *−*0.178, *SE* = 0.051, *p* = 0.002) and higher in the mid-luteal than the early follicular phase (*Estimate* = 0.151, *SE* = 0.051, *p* = 0.009). The progesterone levels during the mid-luteal phase were significantly higher than those during the pre-ovulatory phase (*Estimate* = 116.34, *SE* = 13.05, *p <* 0.001) and early follicular phases (*Estimate* = 138.74, *SE* = 13.05, *p <* 0.001) **(Table 1 and Figure 2)**.

**Table 1:**
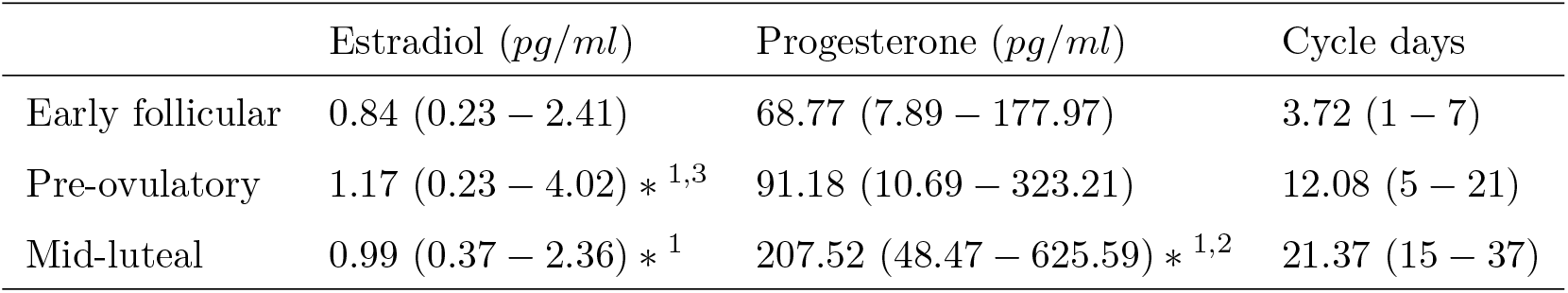
Demographic data for the 60 participants. Significant differences in hormonal levels were observed across phases, with values being significantly higher compared to the early follicular phase (*∗*^1^), pre-ovulatory phase (*∗*^2^), or mid-luteal phase (*∗*^3^). We present the mean values, and their range is in parentheses.

**Figure 2:**
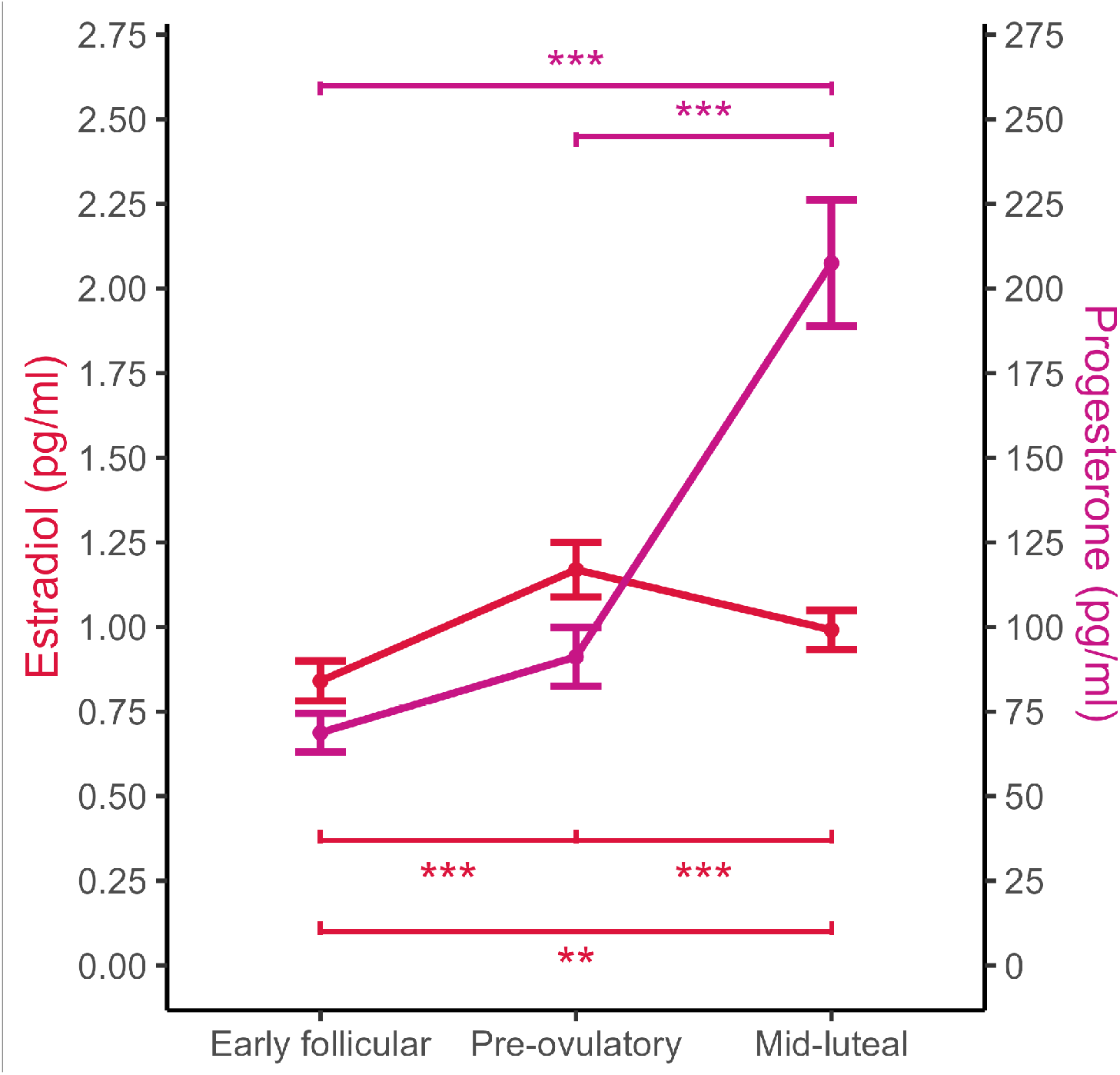
Estradiol (red) and progesterone (magenta) levels during each menstrual cycle phase. Points represent the means, and whiskers are the standard errors. Significant differences are denoted by *** *p <* 0.001, and ** *p <* 0.01.

### Node-metastability across the whole-brain network

We computed the node-metastability measure to study the dynamical complexity underlying the whole-brain functional network in each menstrual cycle phase. We found that the node-metastability was higher in the pre-ovulatory phase compared to the early follicular (FDR-corrected, *p <* 0.001) and mid-luteal (FDR-corrected, *p <* 0.001) phases. Furthermore, we found that the node-metastability was higher in the mid-luteal phase than in the early follicular phase (FDR-corrected, *p <* 0.001) **(Figure 3A)**. In **Figure 3B**, we show the hierarchy for each phase across the whole-brain functional network (i.e., brain areas sorted from highest to lowest node-metastability). The shadow red area represents the 10% brain areas showing the highest node-metastability values for each phase. For the early follicular phase, the brain areas showing the highest values of metastability were primarily located in the attentional networks, DMN, visual, and somatomotor. For the pre-ovulatory phase, brain areas with the highest metastability belonged to the DMN, limbic, subcortical, attentional, and control networks. During the mid-luteal phase, brain areas were mainly located in the subcortical network, DMN, attentional, and control networks. In **Figure 3C**, we show the rendered brains representing the node-metastability for each phase across the whole-brain functional network. It is clear that the pre-ovulatory phase shows the highest metastability values compared to the early follicular and mid-luteal phases.

**Figure 3:**
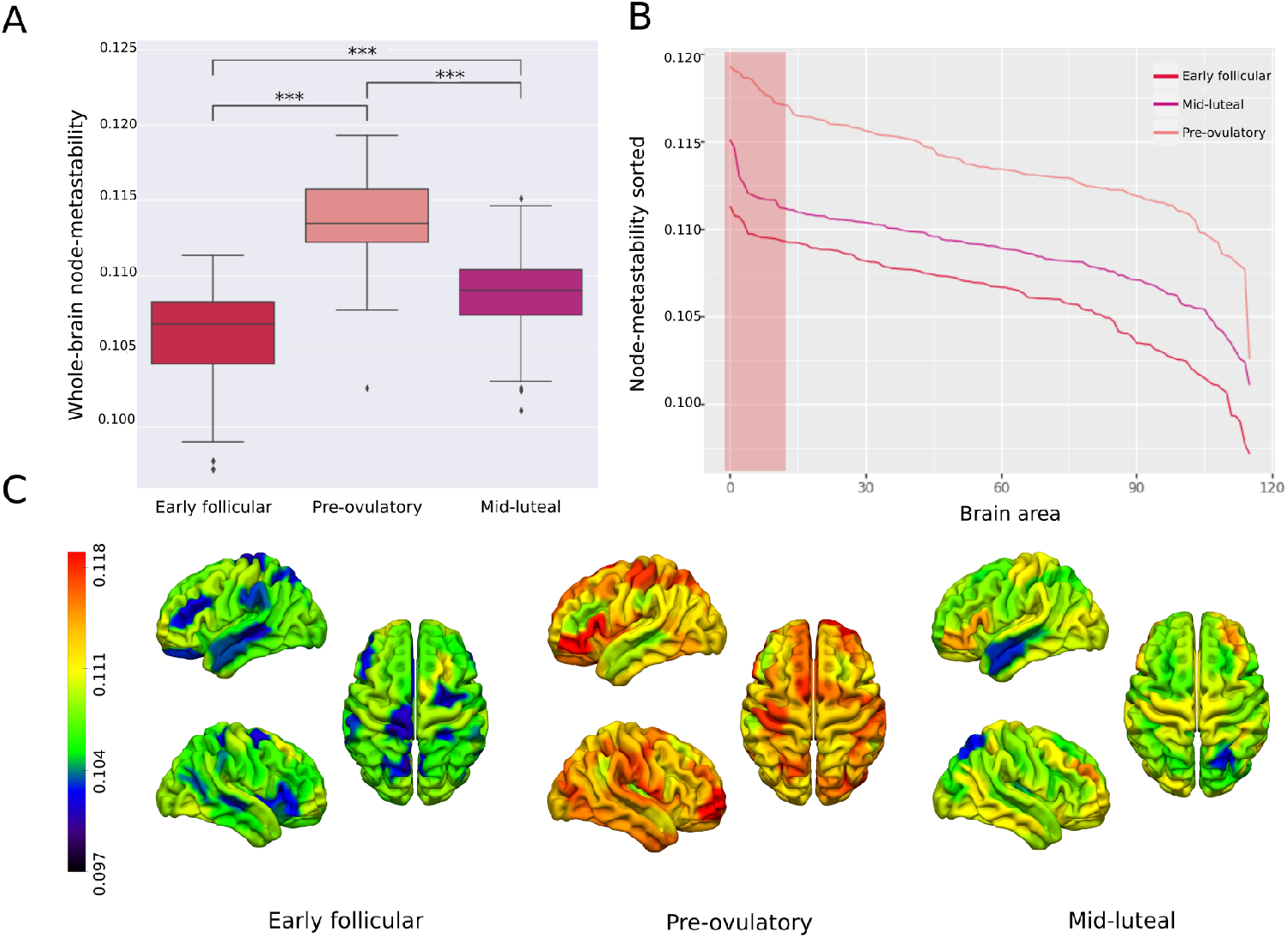
Dynamical complexity of the menstrual cycle phases. **(A)** Node-metastability. The pre-ovulatory phase showed higher node-metastability values across the whole-brain network than the early follicular and mid-luteal phases. P-values are based on Monte–Carlo permutation tests, where *** represents *p <* 0.001. **(B)** Hierarchy. The red area marks the ten regions showing the highest metastability values in each phase. For the early follicular phase, brain areas showing the highest values were primarily located in the salience network, DMN, dorsolateral network, and visual. For the pre-ovulatory phase, the brain areas belonged to the DMN, limbic, subcortical and control networks. During the mid-luteal phase, they were located in the subcortical, DMN, dorsolateral and control networks. **(C)** Renders brains represent the metastability values of the 116 areas for each menstrual cycle phase. The dynamical complexity of the pre-ovulatory phase across the whole-brain networks is clearly more complex than the dynamical complexity of the other two phases.

### Node-metastability across resting state networks

We independently computed the node-metastability for each resting-state network. Differences among menstrual cycle phases for each network are presented in **Figure 4A**. The pre-ovulatory phase showed significantly increased node-metastability compared to the early follicular phase in the DMN, limbic, visual, subcortical networks (FDR-corrected, *p <* 0.001) and salience (FDR-corrected, *p <* 0.05). In contrast, the pre-ovulatory phase showed lower node-metastability compared to the early follicular phase in the dorsolateral attention network (FDR-corrected, *p <* 0.05). Similarly, the mid-luteal phase showed significantly increased node-metastability compared to the early follicular phase in the DMN, limbic, and subcortical networks (FDR-corrected, *p <* 0.001), but showed lower node-metastability in the dorsolateral attention, salience, and somatomotor networks (FDR-corrected, *p <* 0.001). Additionally, compared to the pre-ovulatory phase, the mid-luteal phase showed significantly increased node-metastability only in the DMN network (FDR-corrected, *p <* 0.01) but lower node-metastability in the visual, salience, and somatomotor networks (FDR-corrected, *p <* 0.001). **Figure 4B** shows a radar plot representing the average metastability values for each resting-state network in each menstrual cycle phase.

**Figure 4:**
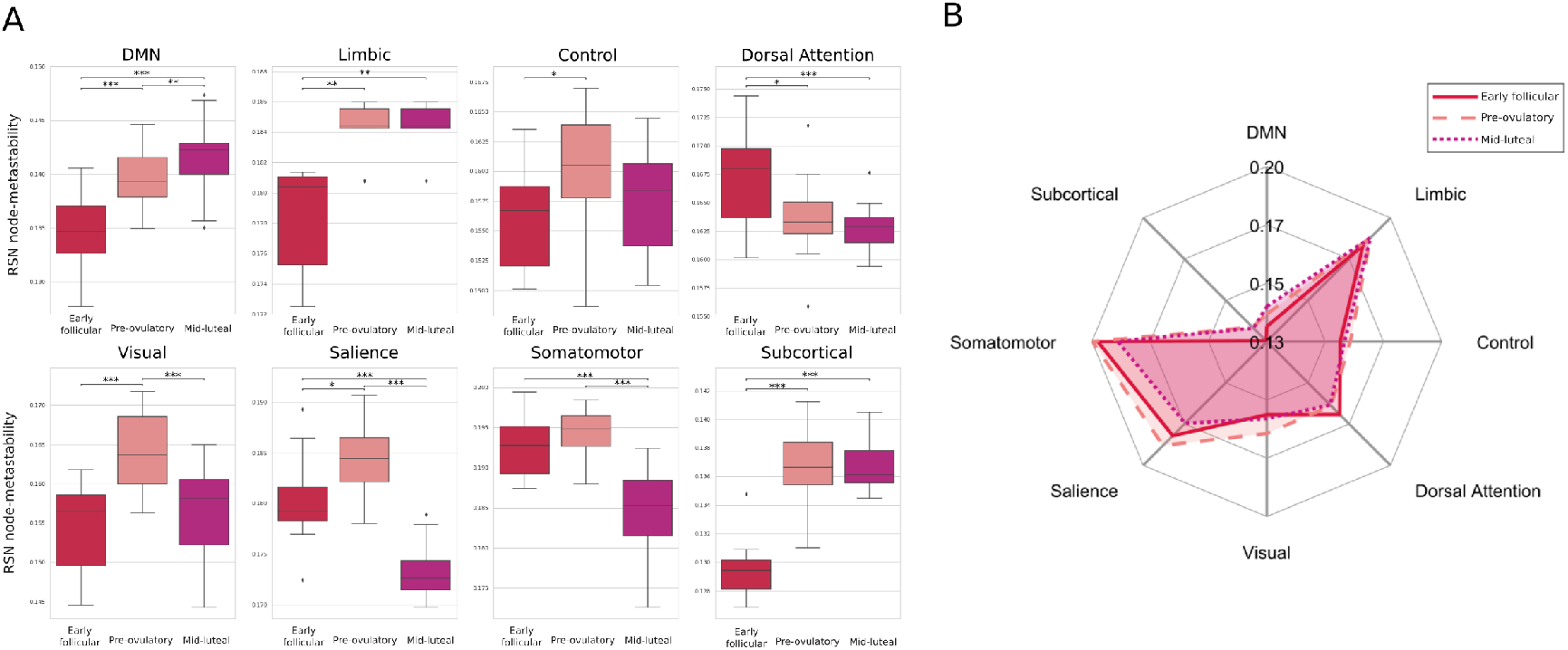
Node-metastability within resting state networks. **(A)** Compared to the early follicular phase, node-metastability was significantly increased in the pre-ovulatory and mid-luteal phases in the DMN, limbic, visual, subcortical, and salience but lower in the dorsolateral network. Compared to the pre-ovulatory phase, the mid-luteal phase showed lower node-metastability in the visual, salience and somatomotor networks and increased node-metastability only in the DMN network. P-values are based on Monte–Carlo permutation tests, where * denotes *p <* 0.05, ** *p <* 0.01 and *** *p <* 0.001. **(B)** The radar plot represents the average metastability values per resting state network for each menstrual cycle phase.

### Multilevel modeling of menstrual cycle effects on brain networks

Mixed effects models for the whole brain showed a significant effect of age on metastability values (*Estimate* = 0.0015, *SE* = 0.00065, *p* = 0.02), indicating higher levels of metastability in the wholebrain network with age. Furthermore, mixed effects models for each resting-state network revealed higher levels of metastability on the control network (*Estimate* = 0.002, *SE* = 0.000, *p* = 0.006) and the dorsolateral attention network (*Estimate* = 0.002, *SE* = 0.000, *p* = 0.0006), with age.

These results suggest that age has a stronger predictive effect on changes in metastability in these networks than levels of hormones. Finally, mixed effects models revealed that progesterone, as well as the interaction between estradiol and progesterone levels, impact node-metastability in the limbic network (progesterone, *Estimate* = 0.000, *SE* = 4.7775*e −* 05, *p* = 0.025; interaction, *Estimate* = *−*7.9082*e −* 05, *SE* = 4.0233*e −* 05, *p* = 0.05), and somatomotor network (progesterone, *Estimate* = 0.002, *SE* = 0.000, *p* = 0.006; interaction, *Estimate* = *−*0.000, *SE* = 5.6642*e −* 05, *p* = 0.027). The results indicate that when progesterone levels are higher (and estradiol levels are lower), the limbic network exhibits greater metastability, while the somatomotor network exhibits reduced metastability.

## Discussion

In this study, we investigated the dynamical complexity underlying three menstrual cycle phases (i.e., early follicular, pre-ovulatory and mid-luteal) in a sample of 60 healthy naturally-cycling women. First, we found that the pre-ovulatory phase showed the highest metastability across the whole-brain functional network, followed by the mid-luteal and early follicular phases. Furthermore, our results revealed that the dynamical complexity of resting-state networks varies according to the menstrual cycle phase. Finally, we found that age impacts the whole-brain functional network, control, and dorsolateral attention networks, as well as progesterone modulates limbic and somatomotor networks.

At the whole-brain network, we found that the dynamical complexity showed the highest variability over time (i.e., higher metastability) during the pre-ovulatory phase, followed by the midluteal, whilst the early follicular showed the lowest **(Figure 3A)**. Higher metastability across the network indicates higher dynamical complexity, implying that the network’s activity exhibits more variability over time (Deco et al., 2017b; Escrichs et al., 2021, 2019). Only a few studies applied a whole-brain dynamic approach and were based on the same dense-sampling single-subject dataset. Specifically, Pritschet et al. (2020) found that estradiol levels impact the DMN and the dorsal attention network, while progesterone was related to reduced coherence across the whole brain. Mueller et al. (2021) showed that brain dynamics were more flexible among prefrontal, limbic, and subcortical nodes during the ovulatory phase (when estradiol levels peak). De Filippi et al. (2021) found that whole-brain turbulent dynamics change between the early follicular and luteal phases. In particular, they found that the luteal phase showed more complex whole-brain turbulent dynamics across long distances in the brain, while the follicular phase was associated with more stable turbulent dynamics. Greenwell et al. (2023) identified recurring network states of high-amplitude dynamic functional connectivity linked to fluctuations in follicle-stimulating and luteinizing hormones. Such studies have shown that the menstrual cycle and hormone concentrations impact whole-brain dynamics. However, they were based on a dense-sample dataset from a young woman in her twenties and presented some limitations. For example, brain activity patterns can vary across women influenced by different factors such as age, hormone sensitivity, or expression of sex hormone receptors (Pritschet et al., 2021). Additionally, the few fMRI time points in the pre-ovulatory phase of this dataset limit statistical power when comparing menstrual cycle phases. Our investigation aligns and extends these findings by examining a large cohort of 60 healthy women during three menstrual cycle phases.

Our findings further reveal a phase-dependent node-metastability across large-scale resting-state networks **(Figure 4)**. In general and in line with previous literature, we found that most networks reached the highest node-metastability during the pre-ovulatory and mid-luteal phases. The only exception was for the dorsal attention network, which exhibited the highest metastability values during the early follicular/menses phase compared to the pre-ovulatory and mid-luteal phases. Specifically, the pre-ovulatory phase showed the highest node-metastability in visual, salience, and somatomotor networks compared to the mid-luteal phase. Most notably, the metastability of the DMN increased significantly across phases (from the follicular to pre-ovulatory to mid-luteal), with the peak occurring during the mid-luteal phase. Additionally, we observed more complex dynamics within limbic and subcortical networks around ovulation and during the mid-luteal phase, which is consistent with prior menstrual cycle resting-state fMRI research (Mueller et al., 2021; Pletzer et al., 2016). Furthermore, our results also align with our previous publications, where we observed distinct brain dynamic patterns within the triple network model (i.e., DMN, salience, and control networks) across menstrual cycle phases (Hidalgo-Lopez et al., 2021), that the DMN exhibits higher levels of turbulent dynamics during the luteal phase (De Filippi et al., 2021), as well as a decrease in the motor network in the mid-luteal phase compared with the early follicular and pre-ovulatory phases (Pletzer et al., 2016). Overall, these results demonstrate menstrual-cycle-related changes in large-scale brain networks.

In order to effectively study the effects of the menstrual cycle on brain function, it is important to use repeated measures designs that account for within-person cycle phase and/or hormone effects (Gehlert et al., 2009; Schmalenberger et al., 2021; Wei et al., 2018). Multilevel modeling is a suitable statistical approach for this purpose as it considers these individual differences as factors while addressing the non-independence of observations resulting from repeated measures data. Therefore, we used a mixed-effects model to examine the effects of age and hormones on node-metastability across the whole-brain and resting state networks. Our results revealed that age significantly increases whole-brain metastability. Furthermore, we observed age-related effects in node-metastability in the control and dorsolateral attention networks. Our results are consistent with previous studies reporting a maximum peak in network efficiency around 40 years old, suggesting an increase in brain dynamics during early adulthood (Cao et al., 2014; Damoiseaux, 2017; Escrichs et al., 2021, 2023; Zhao et al., 2015). Moreover, we found hormone-related changes in node-metastability in the limbic and somatomotor networks. The node-metastability in the limbic network increased with higher progesterone, whereas in the somatomotor network, the node-metastability decreased with greater progesterone. These results are related to previous studies reporting functional connectivity changes in regions of the limbic and somatomotor networks, which are modulated by endogenous progesterone (Arélin et al., 2015).

In summary, our research has important implications for understanding the menstrual cycle’s effects on brain dynamics. This study demonstrates that the menstrual cycle phase affects the dynamical complexity of the whole-brain functional network as well as large-scale resting-state networks. This research may have implications for elucidating the effects of hormones on cognition, mood, and behavior in healthy women and menstrual cycle-related disorders.

## Acknowledgments

A.E. and G.D. were supported by the project eBRAIN-Health - Actionable Multilevel Health Data (id 101058516), funded by EU Horizon Europe. The European Research Council (ERC) Starting Grant 850953 supported B.P. and E.H-L.

## Conflict of interest statement

The authors declare that the research was conducted in the absence of any commercial or financial relationships that could be construed as a potential conflict of interest.

## Notes

### Competing Interest Statement

The authors have declared no competing interest.

